# Postsynaptic burst reactivation of hippocampal neurons enables associative plasticity of temporally discontiguous inputs

**DOI:** 10.1101/2022.06.23.497305

**Authors:** Tanja Fuchsberger, Claudia Clopath, Przemyslaw Jarzebowski, Zuzanna Brzosko, Hongbing Wang, Ole Paulsen

**Affiliations:** Department of Physiology, Development and Neuroscience, Physiological Laboratory, University of Cambridge, Cambridge CB2 3EG, UK; Department of Bioengineering, Imperial College London, South Kensington Campus, London, UK; Department of Physiology, Michigan State University, East Lansing, MI 48824, USA

## Abstract

A fundamental unresolved problem in neuroscience is how the brain associates in memory events that are separated in time. Here we propose that reactivation-induced synaptic plasticity can solve this problem. Previously, we reported that the reinforcement signal dopamine converts hippocampal spike timing-dependent depression into potentiation during continued synaptic activity (Brzosko et al., 2015). Here, we report that postsynaptic bursts in the presence of dopamine produces input-specific LTP in hippocampal synapses 10 minutes after they were primed with coincident pre- and postsynaptic activity. The priming activity sets an NMDAR-dependent silent eligibility trace which, through the cAMP-PKA cascade, is rapidly converted into protein synthesis-dependent synaptic potentiation, mediated by a signaling pathway distinct from that of conventional LTP. Incorporated into a computational model, this synaptic learning rule adds specificity to reinforcement learning by controlling memory allocation and enabling both ‘instructive’ and ‘supervised’ reinforcement learning. We predicted that this mechanism would make reactivated neurons activate more strongly and carry more spatial information than non-reactivated cells, which was confirmed in freely moving mice performing a reward-based navigation task.

## Introduction

For an animal to successfully adjust its behavior to changing environmental demands, it needs to learn to associate a sequence of events or actions to a subsequent outcome, for example a reward, which may be experienced minutes or hours later. How this is achieved in the brain remains unknown. Even the longest time-scale synaptic plasticity event reported so far, behavioral timescale synaptic plasticity (BTSP; Bittner et al., 2017; Milstein et al., 2021), cannot bridge this temporal gap. In machine learning, reinforcement learning algorithms solve this problem by storing a temporary record of the occurrence of an event, known as an eligibility trace, which can later undergo learning changes once the outcome is known (Sutton and Barto, 2018). In biology, dopamine (DA) is thought to represent such a reinforcement learning signal, converting a temporary eligibility trace into a lasting synaptic change (Ljungberg et al., 1992; Schultz et al., 1997; Fremaux and Gerstner, 2016; Gerstner et al., 2018). However, the use of a brief scalar signal to alter the relevant synaptic weights raises two fundamental problems, firstly, that of time scales, and secondly, that of specificity (or credit assignment). To our knowledge, the longest delay reported between a behavioral event and phasic dopamine to successfully reinforce a behavior is a few seconds at most. Yet, in behavioral learning, the time delay between cues and reward may be several minutes or even hours. Moreover, with a longer time delay, the relation between the preceding behavior and the outcome is less clear, questioning what events or actions before the reward should be associated with the outcome. One possible mechanism that could link previous activity with a specific outcome is neuronal reactivation, or replay activity, which in the hippocampus is enhanced by reward (Singer and Frank, 2009; Ambrose et al., 2016).

The rodent hippocampus is important for spatial memories. Spatial representations are built during exploration of an environment, when the hippocampus shows theta activity, and later reactivated or replayed both during sleep (Wilson and McNaughton, 1994; Nadasdy et al., 1999; Lee and Wilson, 2002) and in the awake state (Kudrimoto et al., 1999; Foster and Wilson, 2006; Csicsvari et al., 2007; Diba and Buzsaki, 2007). Reactivation occurs during sharp wave ripples (SWRs), when neurons typically fire action potentials in brief bursts (Buzsaki et al., 1992; Diba and Buzsaki, 2007). It has been reported that stimulation of dopaminergic input promotes reactivation of hippocampal cell assemblies and memory persistence (McNamara et al., 2014).

We investigated the effect of action potential bursts on synaptic plasticity in individual postsynaptic hippocampal CA1 neurons (SWR-associated ‘reactivation’) during dopaminergic modulation (‘reward signal’) after they had undergone a Hebbian pairing protocol (prior exploration-based synaptic ‘priming’). We found that the pairing protocol set an NMDA receptor-dependent silent eligibility trace, which could be converted several minutes later by burst activity in the presence of DA into protein synthesis-dependent long-term potentiation (LTP) mediated by a signaling pathway distinct from that of conventional LTP. Using this synaptic learning rule in a computational model we show that reactivation-induced plasticity increases specificity to reinforcement learning, offering a new mechanism of credit assignment in neural networks. To investigate how reactivation affects the functional properties of neurons *in vivo*, we used chronic calcium imaging of the dorsal CA1 region of the hippocampus in freely moving mice performing a reward-location learning task. We defined neuronal reactivation as activity during immobility at the reward-location in neurons that were already previously active during the reward approach. Neurons that reactivated after finding the reward had increased calcium activity and place map peaks compared to non-reactivated neurons, suggestive of changes in synaptic weights in reactivated neurons.

## Results

### Reactivation during dopaminergic modulation induces LTP

To investigate the effect of burst reactivation of individual postsynaptic CA1 cells during dopaminergic modulation, we monitored the synaptic weights of afferent synapses that had previously undergone a spike timing-dependent synaptic priming protocol. For this we used whole-cell recording of CA1 pyramidal neurons in mouse hippocampal slices (Figure 1A). To be able to distinguish between conventional LTP and reactivation-induced LTP, we used a priming protocol that induces synaptic depression (Andrade-Talavera et al., 2016). Single postsynaptic action potentials followed by presynaptic input (post-before-pre protocol, Δt= -20 ms) led to input-specific synaptic depression (t-LTD; 61% ± 11% vs 100%, t(9) = 3.7, p = 0.005, n = 10; Figure 1B,F). Application of DA alone, without resuming synaptic stimulation after this pairing protocol, did not affect the depression (53% ± 7% vs 100%, t(5) = 7.0, p = 0.0009, n = 6; Figure 1C,F blue trace). Strikingly, postsynaptic action potential bursts (5-6 APs) in the presence of DA, 10 minutes after the pairing protocol, triggered an immediate induction of synaptic potentiation (139% ± 13.6% vs 100%, t(9) = 2.8, p = 0.0199, n = 10; Figure 1D,F red trace). Burst stimulation alone, in the absence of DA, did not prevent synaptic depression (73 ± 10% vs 100%, t(7) = 2.7, p = 0.0321, n = 8; Figure 1E,F black trace). This result suggests that the pairing protocol sets an eligibility trace allowing activated synapses to be selectively altered minutes later by reactivation of the postsynaptic neuron in the presence of DA. To our knowledge, these are the longest-lasting synaptic eligibility traces reported in the brain.

**Figure 1.**
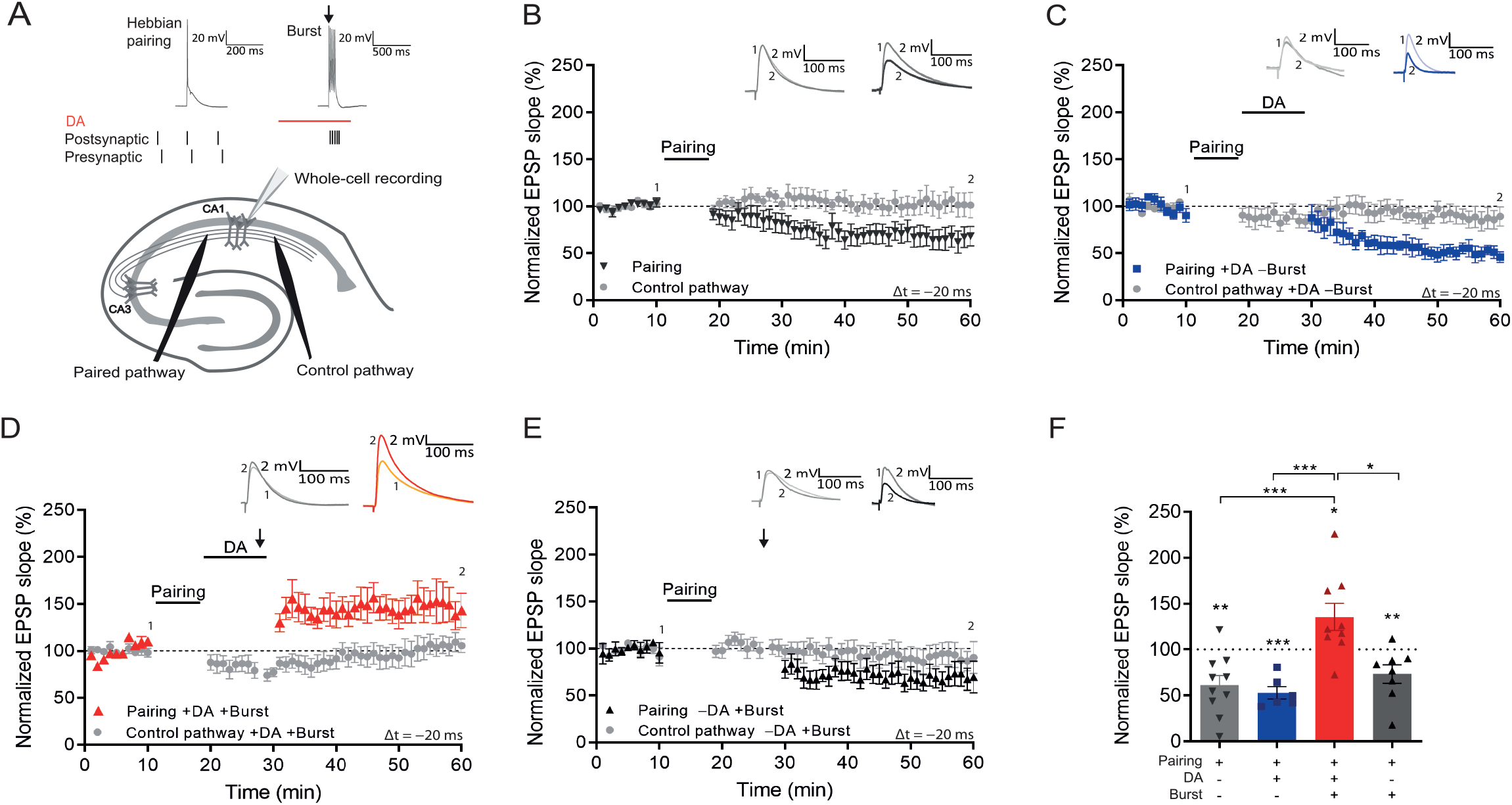
Postsynaptic burst reactivation induces LTP in the presence of dopamine (DA). (A), Schematic of the experimental paradigm (top) and setup (bottom). ↓ = Action potential bursts; Whole cell recording in CA1 stratum pyramidale; electrical stimulation electrodes in stratum radiatum. Plasticity was induced in one pathway (Paired pathway), a second pathway was used for stability control and for confirmation of input specificity (Control pathway). Normalized EPSP slopes were averaged and plotted as a function of time. (B), Post-before pre-pairing protocol leads to input-specific synaptic depression. Pairing protocol (Δt = -20 ms) induces t-LTD (black trace) and does not affect synaptic weights in control pathway (gray trace). (C), DA application after a post-before-pre pairing protocol (Δt = -20 ms) does not prevent t-LTD (+DA – Burst, blue trace) and does not affect synaptic weights in control pathway (gray trace). (D), Application of DA together with action potential bursts of the postsynaptic cell (indicated by black arrow) induces synaptic potentiation after a post-before-pre pairing protocol (Δt = -20 ms) (+DA +Burst, red trace) and does not affect synaptic weights in control pathway (gray trace). (E), The same protocol, without application of DA, leads to synaptic depression (-DA +Burst, black trace) and does not affect synaptic weights in control pathway (gray trace). (F), Summary of results. All traces show an EPSP before (1) and 40 minutes after pairing (2). Plots show averages of normalized EPSP slopes ± s.e.m.

We next investigated the requirements for setting the eligibility trace. First, we found that synaptic potentiation was indeed not observed without prior spike pairing (93% ± 8% vs 100%, t(6) = 0.84, p = 0.43, n = 7; Figure 2A,C). Induction of hippocampal t-LTD requires metabotropic glutamate receptors (mGluRs; Andrade-Talavera et al., 2016). To investigate whether LTD or mGluR signaling is required for burst-induced potentiation, we applied the mGluR antagonist MPEP throughout the recording. This blocked t-LTD (94% ± 6% vs 100%, t(6) = 0.95, p = 0.38, n = 7; Figure 2B,C blue trace; vs control t-LTD 69% ± 7%, n = 7; t(11.5) = 2.67, p = 0.02; Figure 2B,C black trace) but burst-induced potentiation was intact (132% ± 9% vs 100%, t(6) = 3.38, p = 0.015, n = 7 ; Figure 2B,C red trace), suggesting that setting the eligibility trace by Hebbian pairing is distinct from the signaling mechanism that mediates t-LTD.

**Figure 2.**
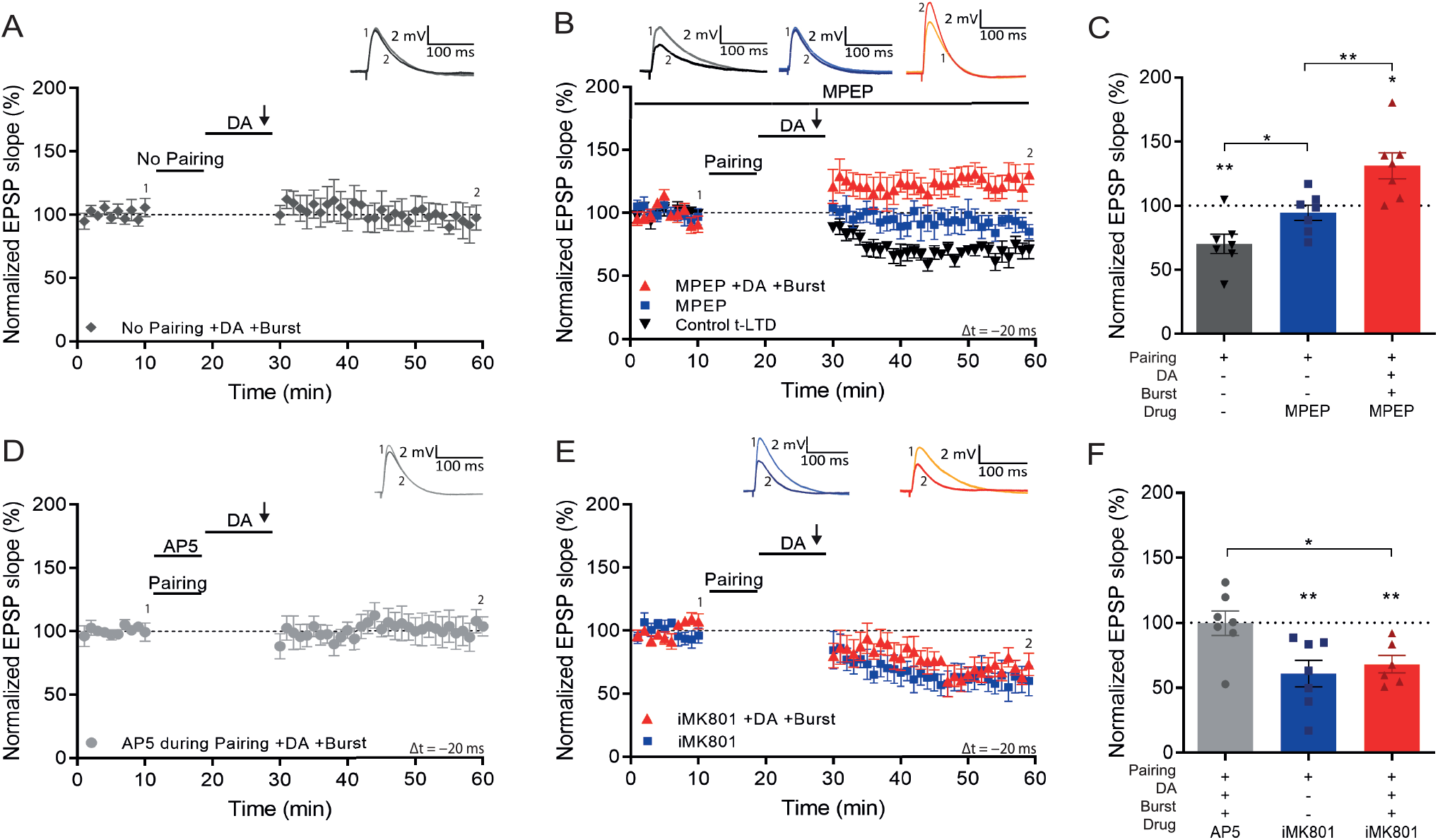
Setting of the eligibility trace is independent of LTD but requires postsynaptic NMDARs. (A), Application of DA and burst stimulation, but without pairing protocol, does not induce potentiation. (B), Post-before-pre pairing protocol (Δt = -20 ms) leads to synaptic depression (t-LTD) (black trace), which is blocked by MPEP (blue trace). MPEP does not block DA- and burst-induced potentiation (red trace). (C), Summary of results. (D), Application of AP5 during pairing blocks DA- and burst-induced potentiation. (E), Postsynaptic intracellular MK801 (iMK801) does not block t-LTD (blue trace) but blocks DA- and burst-induced potentiation (red trace). (F), Summary of results. All traces show an EPSP before (1) and 40 minutes after pairing (2). Plots show averages of normalized EPSP slopes ± s.e.m.

Many forms of hippocampal plasticity require NMDA receptors (NMDARs; Shipton and Paulsen, 2014). We therefore asked if activation of NMDARs during the pairing protocol is required for the synaptic eligibility trace to be set. We found that application of the NMDAR antagonist D-AP5 during pairing (Δt = - 20 ms) abolished both t-LTD and the subsequent burst-induced potentiation, resulting in no change of synaptic weights (100% ± 11% vs 100%, t(5) = 0.006, p = 0.96, n=6; Figure 2D,F). We then investigated whether specifically postsynaptic NMDARs are required for the eligibility trace. Loading the postsynaptic cell with the NMDAR channel blocker MK801 through the recording pipette did not affect t-LTD (61% ± 7% vs 100%, t(5) = 4.77, p = 0.005, n = 7; Figure 2E,F) but completely abolished burst-induced potentiation, leaving synaptic depression instead (61% ± 10% vs 100%, t(6) = 3.85, p = 0.0084, n = 6; Figure 2E,F). These results show a double dissociation of mGluR and non-postsynaptic NMDAR signaling mediating t-LTD, and postsynaptic NMDARs setting an eligibility trace for reactivation-induced potentiation.

We next investigated the mechanism that induces synaptic potentiation during burst stimulation. It was previously shown that activation of NMDARs after pairing is necessary to induce DA-dependent potentiation with subthreshold synaptic stimulation (Brzosko et al., 2015). However, the NMDAR antagonist D-AP5, applied after the pairing protocol but before burst stimulation, did not affect burst-induced potentiation (127% ± 9%, n = 6; t(5) = 2.94, p = 0.032; Figure 3A,C). In contrast, voltage-gated calcium channels (VGCCs) are required as application of nimodipine, an L-type VGCC blocker, before the burst completely abolished LTP and left synaptic depression instead (72% ± 5% vs 100%, t(5) = 5.6, p = 0.0026, n = 6; Figure 3B,C) indicating that, as in some other forms of synaptic potentiation (Grover and Teyler, 1990), Ca^2+^ entry through VGCCs is required for burst reactivation to induce LTP.

**Figure 3.**
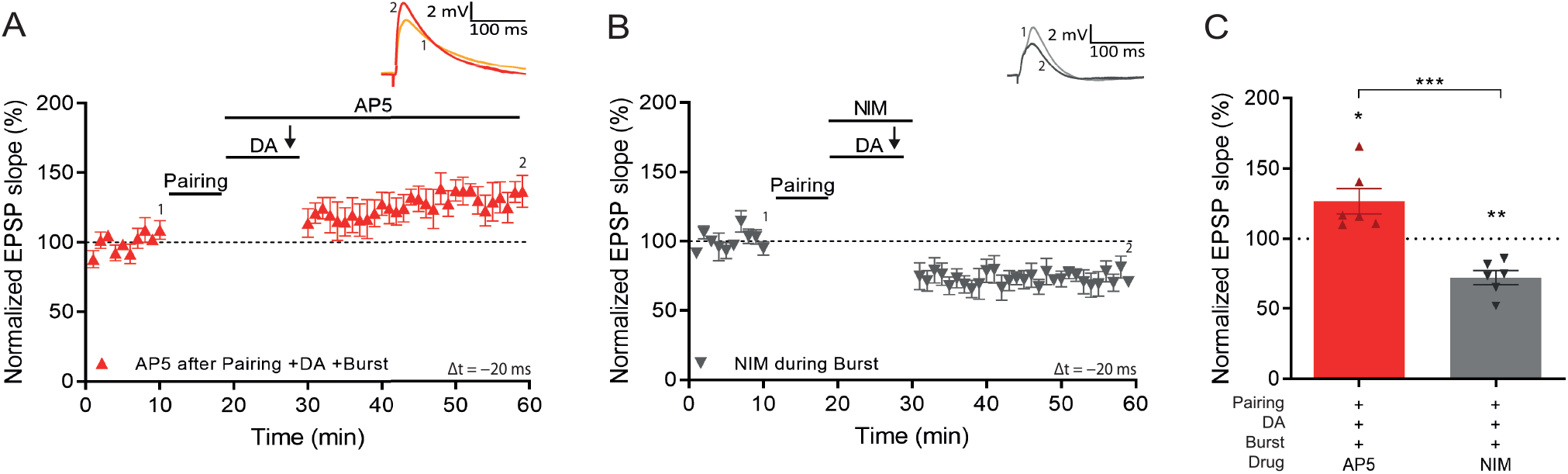
Voltage gated calcium channels mediate DA and reactivation-induced plasticity during burst stimulation. Normalized EPSP slopes were averaged and plotted as a function of time. (A), Application of AP5 after pairing does not prevent DA- and burst-induced potentiation. (B), Application of nimodipine during the burst prevents DA- and burst-induced potentiation leaving synaptic depression. (C), Summary of results. All traces show an EPSP before (1) and 40 minutes after pairing (2). Plots show averages of normalized EPSP slopes ± s.e.m.

### Signaling pathway mediating reactivation-induced LTP

These findings suggest that coincident DA signaling and postsynaptic Ca^2+^ increase enable the potentiation of previously activated synapses. Searching for a potential coincidence detector for DA and intracellular Ca^2+^, we focused on adenylyl cyclases (ACs). They are activated by Gs-coupled dopamine D1/D5 receptor stimulation (Neve et al., 2004), and subtypes AC1 and AC8 are additionally Ca^2+^-stimulated (Wayman et al., 1994; Watson et al., 2000; Ferguson and Storm, 2004). To investigate whether AC subtypes AC1 and/or AC8 are involved in the form of plasticity described here, we tested the induction protocol in AC1/AC8 double knockout (AC DKO) mice (Wong et al., 1999). When postsynaptic burst stimulation in the presence of DA was applied after a negative pairing protocol (Δt = -20 ms; Figure 4Ai) in slices from AC DKO mice, the conversion to potentiation was absent and significantly different from DA- and burst-induced potentiation in slices from wildtype mice (AC DKO, 90% ± 8%, n=6 vs WT, 132% ± 11%, n = 8; t(12) = 2.8, p = 0.015; Figure 4Bi), revealing a role for AC1/AC8 as coincidence detector for DA- and Ca^2+^-induced potentiation. In contrast, conventional, DA-independent t-LTP induced by a pre-before-post pairing protocol (Δt = +10 ms; Figure 4Aii) showed significant potentiation comparable to that seen in wildtype mice (AC DKO, 150% ± 19% vs 100%, t(5) = 2.6, p = 0.049, n = 6; WT, 166% ± 16% vs 100%, t(9) = 4.2, p = 0.0023, n = 10; Figure 4Bii).

**Figure 4.**
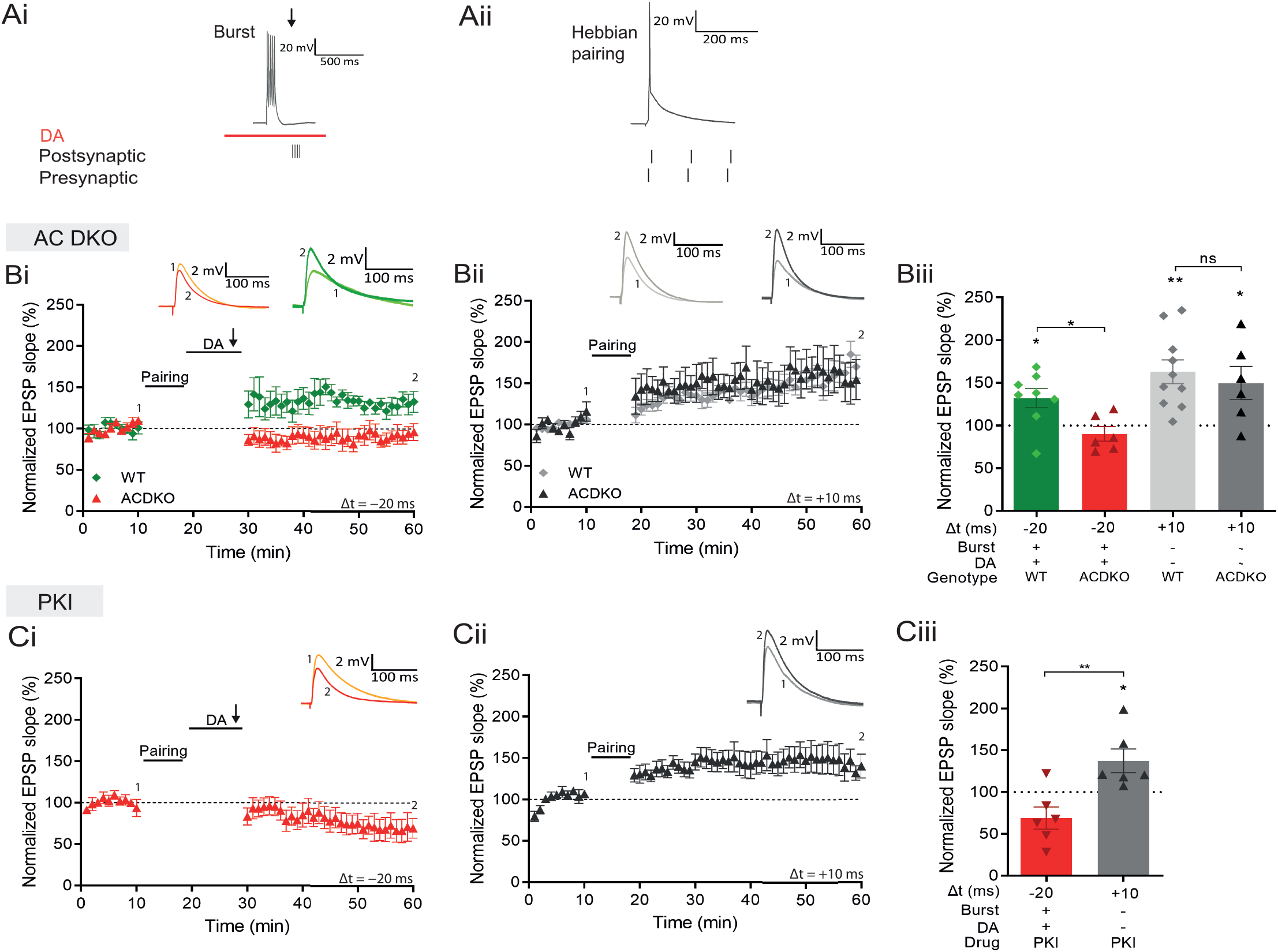
DA and reactivation-induced potentiation requires AC1/AC8 and PKA. Schematics show the difference between the 3 induction protocols. Normalized EPSP slopes were averaged and plotted as a function of time. (Ai), DA and burst stimulation after a post-before-pre pairing protocol (Δt = -20 ms) induces potentiation. (Aii), A pre-before-post pairing protocol induces t-LTP (Δt = +10 ms). (B), AC DKO mice do not show DA-dependent plasticity with postsynaptic bursts (Bi) but shows conventional t-LTP (Bii). Summary of results (Biii). (C), Postsynaptic application of protein kinase inhibitor-(6-22)-amide (PKI) prevents DA-dependent plasticity with postsynaptic bursts (Ci) but leaves conventional t-LTP intact (Cii). Summary of results (Ciii). All traces show an EPSP before (1) and 40 minutes after pairing (2). Plots show averages of normalized EPSP slopes ± s.e.m.

AC activation produces an increase in cyclic adenosine monophosphate (cAMP) which activates protein kinase A (PKA; Sassone-Corsi, 2012). To test whether this signaling cascade is required for reactivation-induced potentiation, we loaded the postsynaptic cell with a PKA blocker, protein kinase inhibitor-(6-22)-amide, through the recording pipette. In this configuration burst stimulation in the presence of DA after the post-before-pre protocol failed to induce synaptic potentiation (69% ± 13% vs 100%, t(5) = 2.4, p = 0.064, n = 6; Figure 4Ci). In contrast, conventional pre-before-post pairing induced significant potentiation, albeit of somewhat reduced magnitude (137% ± 14% vs 100%, t(5) = 2.63, p = 0.0463, n = 6; Figure 4Cii).

The requirement of DA and PKA for burst-induced potentiation is shared with late-phase LTP (Frey et al., 1990; Frey et al., 1993), which requires protein synthesis (Frey et al., 1988). We therefore investigated whether the burst-induced rapid potentiation also requires protein synthesis by delivering the protein synthesis inhibitor anisomycin (AM) to the postsynaptic cell through the recording pipette. We found that, with anisomycin, burst stimulation in the presence of DA no longer induced conversion to potentiation but left a synaptic depression instead, which was significantly different from vehicle control (AM, 52% ± 11% vs 100%, t(5) = 4.5, p = 0.0062, n=6; Figure 5Ai, red trace; vs vehicle, 128% ± 17%, n = 5; t(9) = 3.9, p = 0.0033; Figure 5Ai, green trace). In contrast, conventional t-LTP induced by pre-before-post pairing was unaffected by anisomycin (161% ± 20% vs 100%, t(5) = 3.0, p = 0.029, n = 6; Fig 5Aii, black trace). Furthermore, the post-before-pre pairing protocol induced t-LTD under these conditions (65% ± 6% vs 100%, t(5) = 5.9, p = 0.0019, n = 6; Figure 5Aii, gray trace).

**Figure 5.**
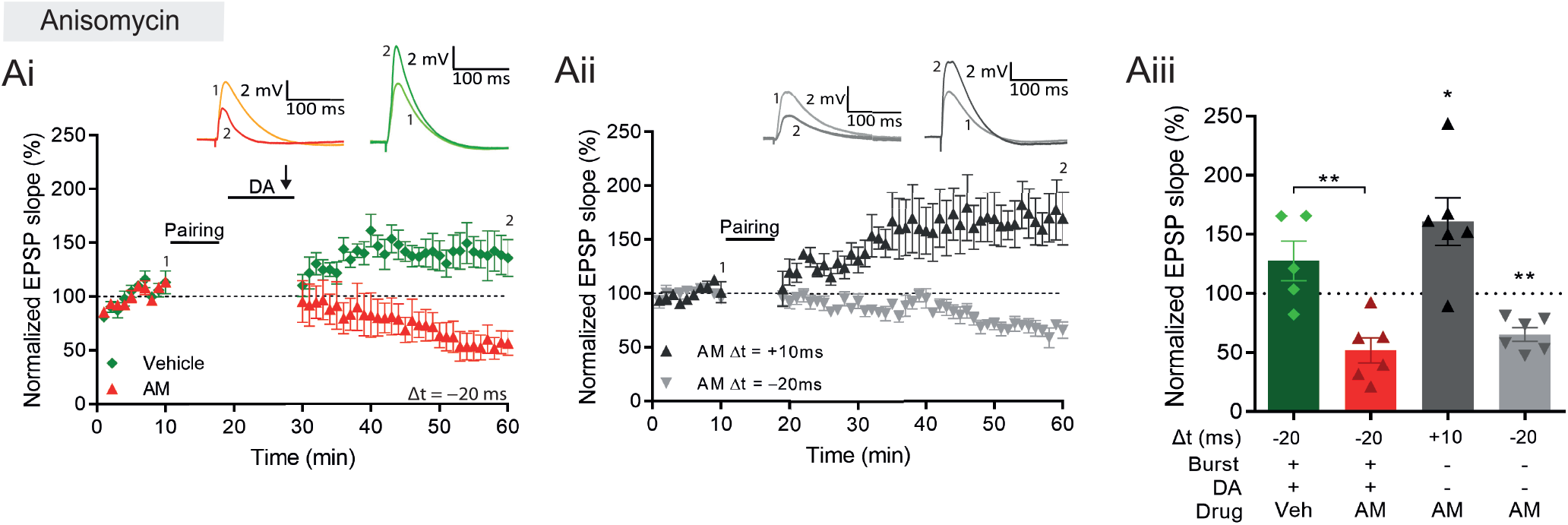
DA and reactivation-induced plasticity requires protein synthesis. (A), Postsynaptic anisomycin prevents DA-dependent plasticity with postsynaptic burst stimulation (Ai), but leaves conventional t-LTD (Aii, gray trace) and t-LTP (Aii, black trace). (Aiii) Summary of results. All traces show an EPSP before (1) and 40 min after (2) pairing. Plots show averages of normalized EPSP slopes ± s.e.m.

### Burst-dependent plasticity increases specificity in reinforcement learning models

These experimental results show that, after a priming event, burst reactivation in the presence of DA induces a rapid form of protein synthesis-dependent LTP. This mechanism would ensure that only salient neuronal activity induces long-term changes in the network. We implemented this synaptic learning rule in a computational model to explore how such plasticity would control learning in a feedforward artificial neural network resembling hippocampal CA1. The learning rule states that the change in synaptic weights *w* between input and output neurons (*i*→*o*) depends on an eligibility trace *e* (set during the initial priming event), the reinforcement signal (dopamine *d*), and reactivation (bursting activity *b*).

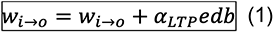

The parameter *αLTP* is the learning rate. When there is no DA or bursts during the trial, the rule is updated to result in depression with a learning rate *αLTD* (see Methods). When using a standard reinforcement learning (RL) rule, which does not depend on burst reactivation, all previously activated synapses are potentiated after receiving the DA signal (Figure 6A). Thus, the global neuromodulatory signal in traditional RL models provides limited information. In contrast, when applying the burst-dependent learning rule (equation 1) to the network, in which potentiation of primed synapses depends on both the reward signal and reactivation, a selected subset of inputs becomes potentiated, while inputs on non-reactivated neurons remain depressed (gray) (Figure 6B). During DA, information is allocated to primed synapses without the need of further coincident pre- and postsynaptic activity, but by reactivation of the postsynaptic neuron alone.

**Figure 6.**
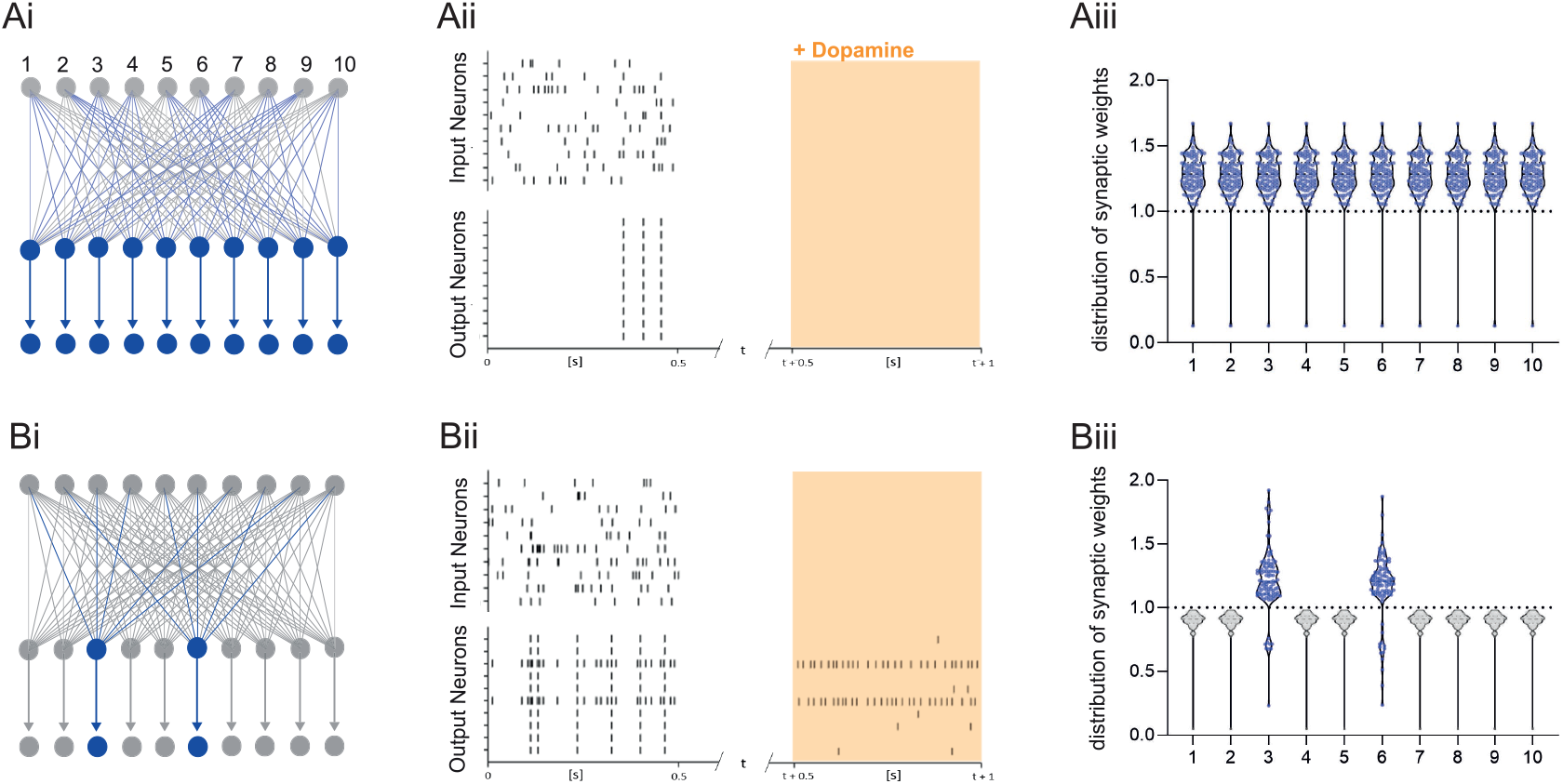
Dopamine-dependent burst-induced plasticity rule reduces the number of neurons in the network encoding a memory. (Ai-iii), Standard reinforcement learning rule shows reward associating inputs 1-10 (blue), leads to potentiation of all synaptic weights (blue). (Bi-iii), Burst-dependent potentiation reduces the number of neurons encoding the memory, leading to potentiation of synapses exclusively onto the most excitable burst-firing neurons 3, 6 (blue).

The broader computational implications of this learning rule depend on the control of postsynaptic neuronal bursting activity. First, it is possible that neurons are recruited to a new memory trace based on their relative neuronal excitability before the training session as suggested by the memory allocation hypothesis (Yiu et al., 2014). According to this scenario, the most excitable cells would be the most likely to show action potential bursts during reactivation and therefore show synaptic potentiation. Alternatively, there is evidence for replay of prioritized experience (Igata et al., 2021), suggesting that cells encoding the most salient events preceding the reward would reactivate, and thereby determine which set of cells would show potentiation during reward (Figure 6B). Assuming prioritized experience reflects experience relevant to the reward, this could help credit assignment in the network. In addition, because of the exclusive requirement of postsynaptic activity for potentiation to occur, this mechanism offers another intriguing possibility, namely that other inputs active at the reward location carries additional information about the nature of the reward, e.g. food, or the reward location, such as the presence of specific landmarks, which could elicit postsynaptic bursting activity. Under this hypothesis neuronal reactivation would serve to associate a specific outcome to the priming event. When different instructive inputs induce bursting each in distinct subsets of neurons during reward, selective increases in synaptic weights would not only enable the encoding of reward, but also distinguish between different rewards (Figure 7A). Finally, we explore how the learning rule performs in a network supervised by feedback synaptic input to strengthen synapses onto specific neurons encoding reward-related features. By allowing feedback input to assign which part of the network is responding, the burst-dependent learning rule enables the network to selectively learn relevant information (Figure 7B), resulting in potentiation of only those synaptic weights (magenta and cyan in Figure 7Biii) when they are active temporally separated, while simultaneous activity leads to less potentiation in those inputs (Figure 7Biv). This provides a mechanism to associate temporally separated, reward-relevant information in a neuronal network. The burst-dependent learning rule provides the network with a gating mechanism for memory allocation to an engram. Thus, burst-induced plasticity reduces the number of neurons encoding the reward location and other reward-related information, increasing the specificity of synaptic memory in a neuronal network.

**Figure 7.**
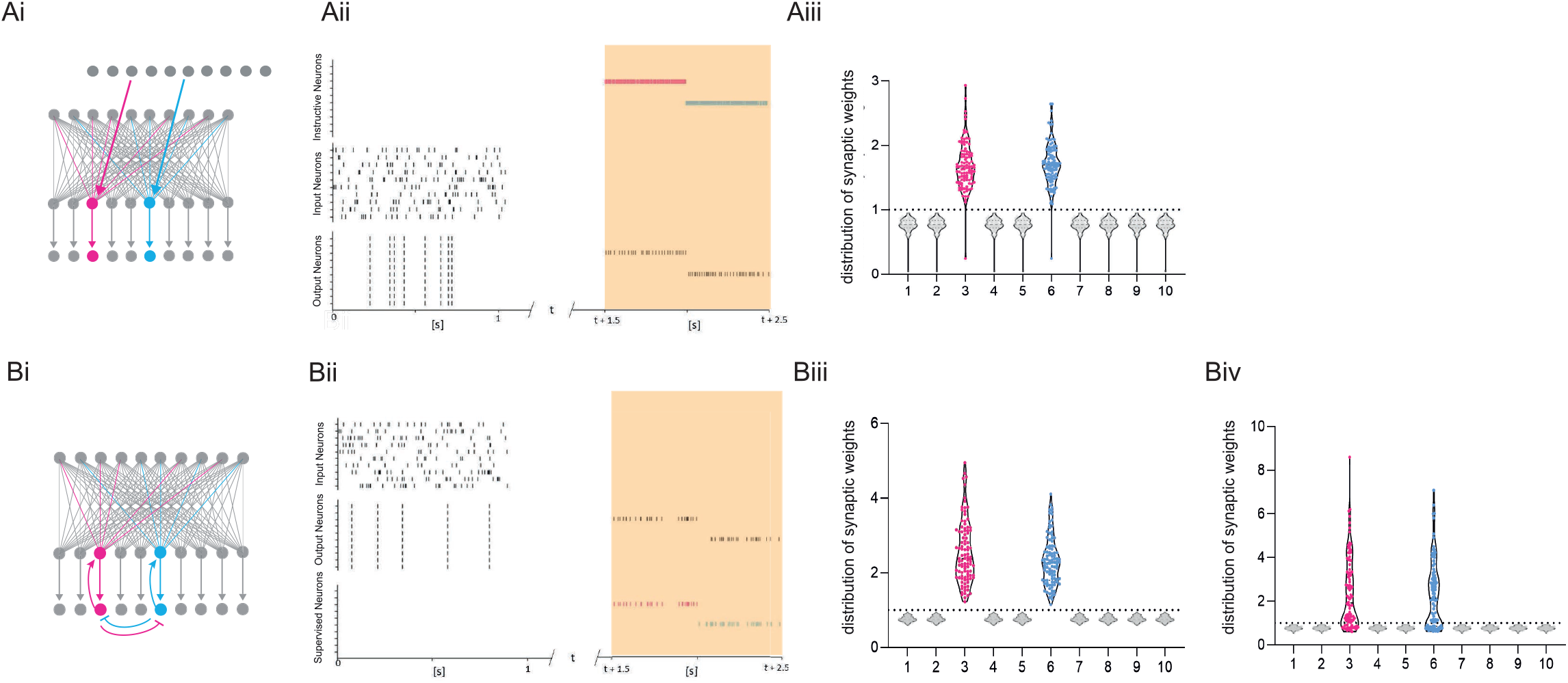
Dopamine-dependent burst-induced plasticity rule enables reinforcement learning (RL) models to encode a specific salient event. (Ai-iii), Instructive RL rule allows two inputs that code for different information to store the memory in separate sets of neurons, thus encoding not only the reward, but also other reward-relevant features 3, 6 (magenta, cyan). (Bi-ii), A supervised network enables burst-eliciting feedback synaptic input to assign credit to select synapses in the network to encode a desired reward identity. (Biii) Time-dependent lateral inhibition at the output neurons supress non-relevant information. When only one of the inputs is active, the animal can learn two different memories over time in neurons 3, 6 (magenta, cyan). (Biv) When both inputs are active at the same time they compete with each other, and neurons 3, 6 (magenta, cyan) are less potentiated.

### Increased calcium responses and spatial information in reactivated CA1 place cells

Based on these results we predicted that, when an animal navigates toward a reward, the previously reactivated hippocampal neurons would be more strongly activated than non-reactivated neurons. To test this prediction, we monitored calcium transients in hippocampal excitatory cells with a head-mounted microscope (Ghosh et al., 2011) while mice foraged on a ‘cheeseboard’ maze with two reward locations, one new to the animal and one previously learnt (Figure 8A). We defined neuronal reactivation as activity during immobility at the reward-location in neurons active during the reward approach. Thus, cells that were active when mice moved towards the reward locations were classified as reactivated if the cells were active again during immobility during and after reward consumption (Foster and Wilson, 2006; O’Neill et al., 2006; Csicsvari et al., 2007) and non-reactivated if no further calcium event was detected after they had reached the reward. Of the cells that were active on the maze in a given trial, 46 ± 1% were reactivated at either or both of the reward locations. There was no detectable difference between the number of cells reactivating at the previously learnt or new reward location (Figure 8B, n = 444 trials).

**Figure 8.**
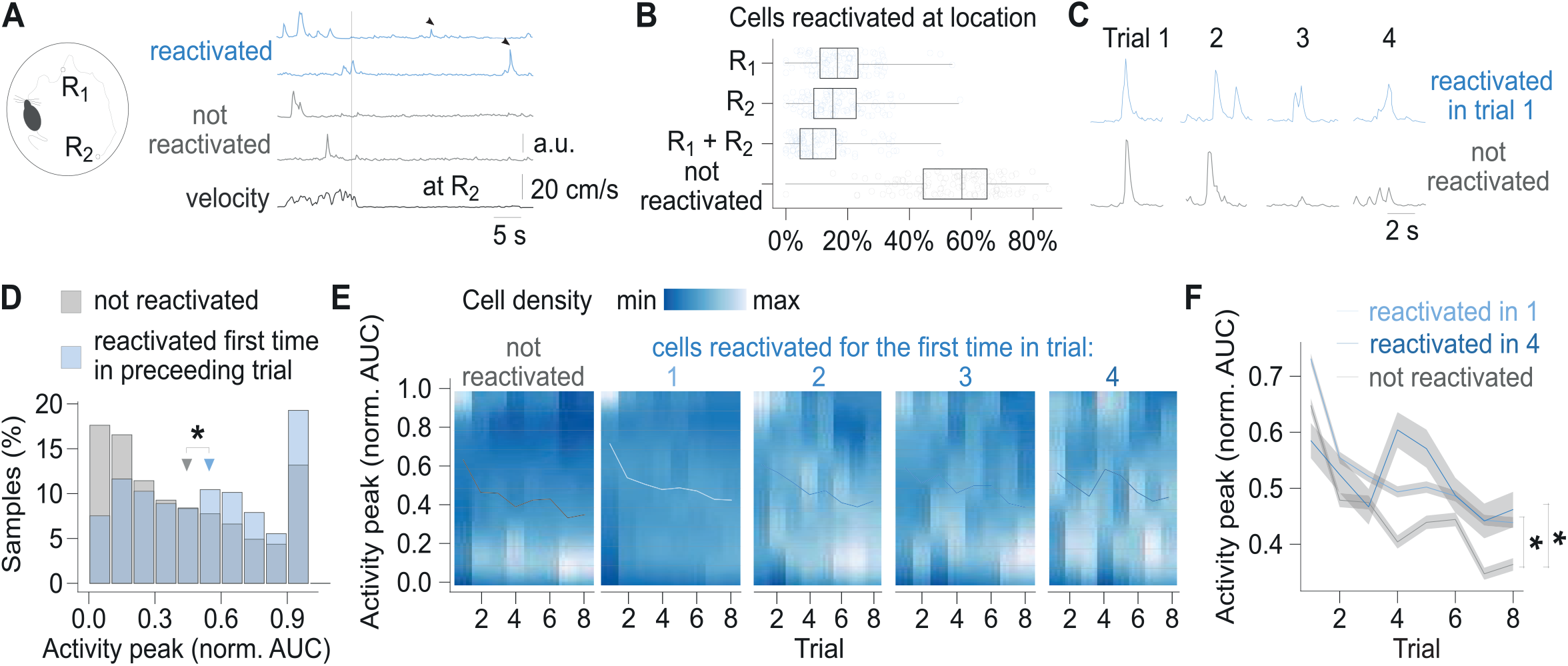
Cells reactivated at reward location have higher activity peaks than non-reactivated cells. (A), Running path of a mouse (left) and calcium traces of example cells (right) in a representative single trial. Mice ran towards two reward locations: one previously learned (R1) and one new (R2). Calcium traces for two cells that reactivated at the reward location and two that did not. Arrows mark reactivations. Vertical line marks time of arrival at R2. (B), Percentage of cells that reactivated at either of the reward locations per trial. Points mark percentage in single trials. Box-and-whisker plots: median, 25 and 75th percentile, and extreme values. (C), Traces centered on activity peaks for two example cells: one that reactivated at the new reward location in trial 1 (top) and one that did not reactivate in any of the trials (bottom). Horizontal dashed lines mark activity peaks. (D), Histogram of activity peaks normalized to maximum AUC value compared between cells that reactivated for the first time in the preceding trial and cells that did not reactivate in any of the trials. Triangles mark mean values. Permutation tests for repeated measures ANOVA: significant effect of trial (F(6, 96) = 4.50, p < 0.001) and reactivation (F(1,16) = 8.55, p = 0.01). n = 6,496 cell trials from 1,404 cells. (E), Density of normalized activity peaks in each trial. Cells are grouped by the trial of their first reactivation. Lines show mean values per trial. (F), Mean calcium peak in each trial for cells that reactivated in trial 1 or trial 4 compared to cells that did not reactivate in any trial. Ribbons extend +/-1 SEM. Cells that reactivated in trial 1 had significantly higher normalized calcium peaks in all trials. Cells reactivated for the first time in trial 4 had significantly higher normalized calcium peaks in trials 4, 5, 7 and 8. Permutation t-tests with Benjamini-Hochberg correction for multiple comparisons and between-animal random effects. n = 6,496 cell trials from 1,404 cells. *p < 0.05.

To compare the strength of neuronal activation, we measured area-under-curve (AUC) of calcium events occurring before and after the reactivation (Figure 8C). Cells with the largest activity peaks during locomotion were the most likely to reactivate at the reward location. Following their first reactivation, they had larger activity peaks than the previously non-reactivated cells (Figure 8D). Similarly, of the two cell groups with matching event rates in a given trial, the one whose cells were reactivated, had larger peaks in the trial following the first reactivation than the one whose cells did not reactivate (Figure 8-S1A). Typically, the calcium activity peak during locomotion increased in the trial immediately before the first reactivation and remained higher than the activity peaks in cells that were not reactivated throughout the day (Figure 8E,F), suggesting that reactivated cells have undergone a lasting change. Photobleaching could affect the magnitude of the detected calcium signal. As the reactivated cells were more active than non-reactivated cells (Figure 8-S1A), we would expect them to be more affected by photobleaching, but nevertheless, the magnitude of their calcium transients remained higher than in the non-reactivated cells. 60% of the reactivated cells (n = 971 from 7 mice) and 42% of the non-reactivated cells (n = 284 from 7 mice) showed activity at specific locations and were classified as place cells (see Methods). Both place cells and non-place cells showed higher calcium peaks following reactivation and the increase in place cells was not significantly different from that in non-place cells (Figure 8-S1B,C).

To investigate any learning-induced changes in place maps we assessed how the location-averaged activity in place cells changed with reactivation from the first to the second half of the trials (Figure 8-S2A). The place map peaks increased significantly more in the place cells that reactivated in trial 4 or later than in the non-reactivated place cells (Figure 8-S2B, mean place map peak ratio 1.4 ± 0.1 vs 1.1 ± 0.1, n = 194 reactivated and n = 284 non-reactivated place cells; permutation test for repeated measures ANOVA: p = 0.009). The change in spatial information also significantly differed between the two groups (Figure 8-S2C, mean change by 0.014 ± 0.015 vs -0.046 ± 0.014 a.u.; permutation test for repeated measures ANOVA: p = 0.005), but there was no significant difference in place map stability (Figure 8-S2D, mean correlation of 0.45 ± 0.02 vs 0.47 ± 0.02; permutation test for repeated measures ANOVA: p = 0.59). Reactivated place cells conveyed more spatial information in late trials compared to place cells that did not reactivate (0.14 ± 0.01 vs 0.10 ± 0.01 a.u.; permutation test for repeated measures ANOVA: p = 0.07). This learning-associated increase in calcium response and spatial information supports a reactivation-dependent LTP-like mechanism *in vivo* (Cacucci et al., 2007).

## Discussion

In summary, we investigated the effects of postsynaptic neuronal reactivation on hippocampal synaptic plasticity, reinforcement learning, and spatial coding. We found that postsynaptic burst reactivation of CA1 pyramidal cells in the presence of the reward signal DA rapidly potentiates synapses that have previously undergone a spike timing-dependent priming protocol, providing direct evidence for reactivation-induced synaptic plasticity. A computational model showed how this learning rule increases specificity in reinforcement learning models and recording from freely moving mice showed that neurons that reactivated at reward locations had enhanced CA1 place cell calcium signals and carried more spatial information than cells that did not reactivate.

The results suggest that reactivation-induced plasticity is mediated by two sequential coincidence detectors: postsynaptic NMDARs detecting coincident pre- and postsynaptic activity and AC1/AC8 as coincidence detector of DA and reactivation-evoked Ca^2+^ increase. Although we used a Hebbian protocol, we cannot exclude the possibility that activation of postsynaptic NMDA receptors without postsynaptic action potentials would be sufficient to set the eligibility trace. The signaling cascade leading to synaptic potentiation involves postsynaptic PKA and protein synthesis, which are not required for conventional early LTP (Park et al., 2014). Traditionally, LTP has been classified into early- and late-phase LTP (Frey et al., 1988; 1990), and it has been reported that dopaminergic signaling is required for maintenance of late-phase LTP (Frey et al., 1990; Huang and Kandel, 1995; Matthies et al., 1997). The DA-dependent form of plasticity we describe here shares properties with ‘late-phase’ LTP, including a role of postsynaptic action potentials during induction (Dudek and Fields, 2002) and a requirement of protein synthesis for expression (Frey et al., 1988; Huang and Kandel, 1995). However, it is remarkably fast, suggesting a dissociation between different signaling pathways, rather than different temporal phases of LTP. The mechanistic basis for the surprising involvement of protein synthesis in this rapidly induced form of plasticity remains to be investigated.

Our experimental findings uncover a synaptic learning rule that could support a two-stage model of hippocampal memory formation (Buzsaki, 1989), in which eligibility traces are laid down during hippocampal theta activity, with subsequent postsynaptic burst reactivation during sharp wave-ripples inducing LTP at those synapses. Exploring possible computational implications of this synaptic learning rule, we first tested it with a neural network in which the most excitable cells are reactivated. The memory allocation hypothesis suggests that learning triggers a temporary increase in neuronal excitability, enabling the linking of individual memories acquired close in time (Silva et al., 2009; Cai et al., 2016; Sehgal et al., 2018). We found that our learning rule selectively strengthens the reactivated synapses, linking together the memory allocation hypothesis with burst reactivation-dependent plasticity. Interestingly, it was recently reported that DA released by LC cells projecting to dCA1 have a key permissive role in contextual memory linking (Chowdhury et al., 2021). Moreover, the rule could also accommodate a temporally discontiguous instructive learning signal or a specific supervisory feedback signal. Thus, it is possible that DA serves as a scalar reward signal which combines with a vectorial representation of the reward identity which triggers the reactivation of a specific subset of neurons. Thus, it was suggested that the direct pathway from the entorhinal cortex could provide an instructive signal to generate accumulation of CA1 place cells at the reward location (Grienberger and Magee, 2021). An attractive possibility is that a downstream subset of neurons active during navigation serves as an instructive input onto upstream neurons during reward. This would be consistent with a dual role of DA as reinforcement signal and enhancer of reverse replay (Ambrose et al., 2016), establishing a predictive chain of potentiated synapses towards the rewarded outcome reminiscent of the successor representation (Dayan, 1993; Stachenfeld et al., 2017). It was suggested in a computational study that feedback regulation of synaptic plasticity by bursts in higher hierarchical circuits can coordinate lower-level connections (Payeur et al., 2021). Our results reveal a possible biological substrate to support such a mechanism and re-emphasize the importance of bursting activity for synaptic plasticity (Lisman, 1997; Pike et al., 1999).

The hippocampus receives dopaminergic input from two main sources, the ventral tegmental area (Scatton et al., 1980; Gasbarri et al., 1994), signaling reward, and the locus coeruleus (Smith and Greene, 2012; Kempadoo et al., 2016), thought to signal novelty (Takeuchi et al., 2016; Wagatsuma et al., 2018), but more recently implicated also in spatial reward learning (Kaufman et al., 2020). In a reward location learning task, we found that calcium signals during locomotion were higher in CA1 principal cells that reactivated at reward location. The signal increased in the trial preceding the first reactivation, following which the signal remained higher in all subsequent trials. Moreover, place cells that reactivated showed higher spatial information than non-reactivating place cells. The overall stability of calcium signals and spatial information is broadly consistent with earlier reports using one-photon imaging in freely moving mice (Ziv et al., 2013). Although it was recently reported in head-fixed mice that reactivation increases long-term stability of place cells that have fields distant from the reward after several days (Grosmark et al., 2021), we did not see a significant increase in place cell stability after a single reactivation of the cell at the reward location. Irrespectively, the difference in both calcium signal and spatial information in reactivating *vs* non-reactivating cells is suggestive of plasticity related to reactivation events. More work will be required to establish under what conditions the novel burst-induced potentiation mechanism is engaged during hippocampus-dependent learning and memory.

## Author Contributions

T.F. and O.P. designed the experiments. T.F., P.J. and Z.B. conducted the experiments and analyzed the data. C.C. developed the computational model. H.W. provided transgenic AC DKO mice. T.F., C.C., P.J. and O.P. wrote the manuscript.

## Acknowledgements

This research was supported by the Biotechnology and Biological Sciences Research Council, U.K. We are grateful for discussions of this project with other members of the Neuronal Oscillations Group.

## Competing Interests statement

The authors declare no competing interests.

## METHODS

### Mice

Experimental procedures and animal use were performed in accordance with UK Home Office regulations of the UK Animals (Scientific Procedures) Act 1986 and Amendment Regulations 2012, following ethical review by the University of Cambridge Animal Welfare and Ethical Review Body (AWERB). All animal procedures were authorized under Personal and Project licences held by the authors.

Mice were housed on a 12-hr light/dark cycle at 19–23 °C and were provided with food and water *ad libitum*. Experiments were carried out on wild-type C57/BL6J mice (Harlan, Bicester, UK or Central Animal Facility, Physiological Laboratory, Cambridge University), and adenylate cyclase double knock-out (AC DKO) mice, which have the genes for both AC1 and AC8 deleted globally. This mouse line was generated as described previously^25^ and was imported from Michigan State University, MI, US. For *in vivo* experiments, 7 adult male Thy1 – GCaMP6f transgenic mice were used (Dana et al., 2014) (Jax: 024276). Mice were housed with 2-4 cage-mates in cages with running wheels.

### Electrophysiology

#### Slice preparation

Mice of both sexes at postnatal day (P) 12-19 were used in this study. Mice were anesthetized with isoflurane (4% isoflurane in oxygen) and decapitated. The brain was rapidly removed and immersed in ice-cold artificial cerebrospinal fluid (ACSF) containing (in mM): 126 NaCl, 3 KCl, 26.4 NaH2CO3, 1.25 NaH2PO4, 2 MgSO4, 2 CaCl2, and 10 glucose (pH 7.2, 270–290 mOsm/L). The ACSF solution was continuously bubbled with carbogen gas (95% O2, 5% CO2). Horizontal slices (350 μm thick) were sectioned with a vibrating microtome (Leica VT 1200S, Leica Biosystems, Wetzlar, Germany). The slices were then incubated for at least 60 min in ACSF at room temperature in a submerged-style storage chamber before recording. Slices were used for 1–7 hr following sectioning.

#### Whole-cell patch clamp recording

For recordings, individual slices were transferred to an immersion-type recording chamber and perfused with ACSF (2 ml/min) at 24–26 °C. Neurons were visualized and selected using infrared differential interference contrast (DIC) microscopy using a 40X water-immersion objective. The hippocampal subfields were visually identified and whole-cell patch-clamp recordings were performed on CA1 pyramidal neurons. For stimulation of Schaffer collaterals, monopolar stimulation electrodes were placed in stratum radiatum. Patch pipettes (4–7 MΩ) were pulled from borosilicate glass capillaries (0.68 mm inner diameter, 1.2 mm outer diameter) using a P-97 Flaming/Brown micropipette puller (Sutter Instruments Co., Novato, California, USA). Pipettes were filled with a solution containing (mM): 110 potassium gluconate, 4 NaCl, 40 HEPES, 2 ATP-Mg, 0.3 GTP (pH 7.2–7.3, 270–285 mOsm/L). The liquid junction potential was not corrected for.

All experiments were performed in current-clamp mode. Cells were accepted for the experiment if their resting membrane potential was between −55 and −70 mV. The membrane potential was held at −70 mV throughout the recording by direct current application via the recording electrode. Before the start of each recording, all cells were tested for regular spiking responses to positive current steps—characteristic of pyramidal neurons.

#### Stimulation protocol

Excitatory postsynaptic potentials (EPSPs) were evoked alternately in two input pathways (test and control) by direct current pulses at 0.2 Hz (stimulus duration 50 μs) through metal stimulation electrodes. The stimulation intensity was adjusted (100 μA– 500 µA) to evoke an EPSP with peak amplitude between 3 and 8 mV. After a stable EPSP baseline period of at least 10 min, STDP was induced in the test pathway by repeated pairings of single evoked EPSPs and single postsynaptic action potential elicited with the minimum somatic current pulse (1–1.8 nA, 3 ms) via the recording electrode. Pairings were repeated 100 times at 0.2 Hz. Spike-timing intervals (Δt in ms) were measured between the onset of the EPSP and the onset of the action potential.

Alternate stimulation of EPSPs was resumed immediately after the pairing protocol and monitored for at least 40 min. For the burst stimulation protocol, stimulation of EPSPs was not resumed for an additional 10 mins and at the end of that period, 5-6 bursts of action potentials were elicited with an inter-burst interval of 0.1 Hz, by somatic current pulses via the recording electrode (5x 1.8 nA, 10 ms each). Immediately after the bursts, stimulation of EPSPs was resumed and monitored for at least 30 min. Presynaptic stimulation frequency to evoke EPSPs remained constant throughout the experiment. The unpaired pathway served to verify input-specificity and as a stability control. The burst stimulation protocol is summarized in Figure 1a (top).

#### Drugs

Drugs were bath-applied to the whole slice through the perfusion system by dilution of concentrated stock solutions (prepared in water or DMSO) in ACSF, or by adding the drugs to the patch pipette solution when it was applied intracellularly to the postsynaptic cell only. If the drug was not water-soluble, vehicle control experiments were carried out. For each set of recordings, interleaved control and drug conditions were carried out and were pseudorandomly chosen. The following drugs were used in this study: 100 μM dopamine hydrochloride (Sigma–Aldrich, Dorset, United Kingdom), 100 μM D-AP5 (Tocris Bioscience, Bristol, United Kingdom), 10 µM nimodipine (Tocris Bioscience), 50 μM forskolin (Tocris Bioscience), 1 µM PKA inhibitor fragment (6-22) amide (Tocris Bioscience), 0.5 mM anisomycin (stock solution in EtOH; Tocris Bioscience) and 20 µM NASPM (Tocris Bioscience), 1 mM MK801 (Tocris Bioscience).

#### Data acquisition and data analysis of slice recordings

Data were collected using an Axon Multiclamp 700B amplifier (Molecular Devices, Sunnyvale, California, USA), acquired and digitized at 5 kHz using an Instrutech ITC-18 A/D interface board (Instrutech, Port Washington, New York, USA) and custom-made acquisition procedures in Igor Pro (WaveMetrics, Lake Oswego, Oregon, USA).

All experiments were carried out in current clamp (‘bridge’) mode. Series resistance was monitored (10–15 MΩ) and compensated for by adjusting the bridge balance. Data were discarded if series resistance changed by more than 30%. Offline analyses were done using custom-made procedures in Igor Pro. EPSP slopes were measured on the rising phase of the EPSP as a linear fit between the time points corresponding to 25–30% and 70–75% of the peak amplitude. For statistical analysis, the mean EPSP slope per minute of the recording was calculated from 12 consecutive sweeps and normalized to the baseline. Normalized ESPS slopes from the last 5 min of the baseline (immediately before pairing) and from the last 5 min of the recording were averaged. The magnitude of plasticity, as an indicator of change in synaptic weights, was defined as the average EPSP slope after pairing expressed as a percentage of the average EPSP slope during baseline.

#### Statistical analysis of slice recordings

Statistical comparisons were performed using one-sample two-tailed, paired two-tailed, or unpaired two-tailed Student’s *t*-test, with a significance level of α = 0.05. Data are presented as mean ± s.e.m. Significance levels are indicated by *p < 0.05, **p < 0.01, ***p < 0.001. All datasets passed the test for normality using the Shapiro-Wilk test (α = 0.05).

### Computational Modeling

We simulated a set of n = 10 output neurons, which each received input from 10 input neurons. When an instructive input was added, output neurons additionally received input from two out of 10 instructive neurons. Each output neuron projected uniquely onto one readout neuron which again projected back to the output neuron in a one-to-one mapping. Each output neuron was modelled as an Integrate-and-Fire neuron and 100 trials were simulated, where the voltage v is described by

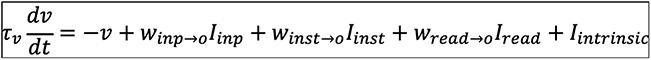

where *τ_ν_* = 10*ms* is the membrane time constant, *I_inp_* are the spike trains of the input neurons, *I_inst_* are the spike trains from the instructive neurons, *I_read_* are the spike trains from the read-out neurons, *w_inp_*_→*o*_ are the weights from the input to the output neurons, *w_inst_*_→*o*_ = 1/*N* are the weights from the instructive neurons to the output neurons, and w_read→*o*_ = 1 are the weights from the readout to the output neurons. In addition, each neuron was receiving an intrinsic current *I_intrinsic_* = *α_intrinsic_η*, where *η* is simply a white noise drawn from a uniform distribution between 0 and 1 and *α_intrinsic_* is the magnitude of the current (*α_intrinsic_* is taken as 0 unless otherwise stated). When the voltage crosses the firing threshold = 0.4, the neuron is reset to the resting potential v = 0. The read-out neurons were also modeled as an Integrate-and-Fire neuron

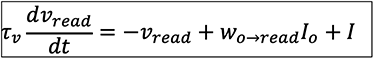

where *I_o_* are the spike-trains of the output neurons, *w_o_*_→*read*_ = 1/*N* are the weights from the output to the readout neurons and *I* are the spike trains from the supervised neurons. Similarly, if the voltage crossed the firing threshold = 0.4, the neurons emit a spike and the voltage is reset to 0. *w_inp_*_→o_ were plastic under the following rule. Every time the input neurons are firing (at *t^pre^*), they are leaving a trace *x^pre^*, 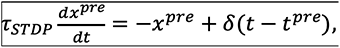 where *τ_STDP_* = 10*ms* is the trace time constant. Similarly, every time the output neuron spikes (at *t^post^*), it is leaving a trace *_xpost_*, 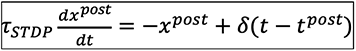.

The eligibility trace e is described as

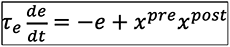

where *τ_e_* = 10 min is the eligibility time constant. The *w_inp→o_* are potentiated if there is an eligibility trace together with dopamine and a postsynaptic burst:

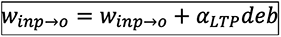

where *α_LTP_* = 0.0002 is the learning rate, d = 0 if there is no dopamine, and d = 1 if there is dopamine. The postsynaptic burst b is detected as follows. We first computed a trace *x^burst^* as

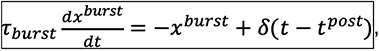

where *τ_burst_* = 5*ms* is the burst time constant. We set b = 1 if *x^burst^* > *burst_threshold_*, to 0 otherwise, with *burst_threshold_* = 1.1. If there was no dopamine nor bursts during the whole trial, then the updated rule resulted in a depression

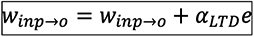

where *α_LTD_* = *α_LTP_*/500 is the learning rate for the depression. Weights are bound to stay positive. The weights *w_inp→o_* are initialized to 1/N. We simulated the network for 2s in Figure 6 and 2.5s in Figure 7. The input neurons were firing Poisson statistics at 20 Hz for the first 500 ms of the trial (pairing phase). Dopamine was present during the last 500 ms of the trials in Figure 6 and the last 1 s in Figure 7. The network was simulated for 100 trials (error bars are standard deviations).

In Figure 6A, a standard reinforcement learning rule was used: *w_inp→o_*= *w_inp→o_* + *α_RL_de*), where *α_RL_* = *α_LTP_*/50. In Figure 6B, a burst dependent rule was used. Output neurons 3 and 6 had an increased excitability. We modeled that by decreasing the firing threshold to 0.25. All neurons also received an intrinsic current with *α_intrinsic_* = 0.06 during the last 500 ms of the trail. In Figure 7A, the instructive neuron 3 fired Poisson statistics at 500 Hz from time 1.5-2 s and the instructive neuron 6 fired Poisson statistics at 500 Hz from time 2-2.5 s. In Figure 7B ii/iii, the read-out neuron 3 received additional Poisson inputs at 60 Hz from time 1.5-2 s and the read-out neuron 6 from time 2-2.5 s. For Figure 7B iv however, the read-out neurons 3 and 6 received their additional Poisson inputs at the same time, from 1.5-2.5 s. In Figure 7B, we added lateral inhibition, where each read-out neuron received an inhibitory filtered version (with a time constant of 50 ms) of the spike trains of other read-out neurons, with weights of -1/N (without self-connection).

### Behavior

#### Surgery

Mice underwent two surgeries: the first one to implant a GRIN lens directly above the cells of interest, and the other to fix an aluminum baseplate above the GRIN lens for later attachment of the miniature microscope. The procedures followed the protocol described before by Resendez et al. (2016). Surgeries were carried out following minimal standard for aseptic surgery. Analgesic (Meloxicam, 2 mg.kg-1 intraperitoneal) was administered 30 min prior to surgery. Mice were anesthetized with isoflurane (5% induction, 1-2% maintenance, Abbott Ltd, Maidenhead, UK) mixed with oxygen as carrier gas (flow rate 1.0-2.0 L.min-1) and placed in a stereotaxic frame (David Kopf Instruments, Tujunga, CA, USA). The skull was exposed by making skin incision and Bregma and Lambda were aligned horizontally. A 1.5–2 mm-wide craniotomy was drilled above the implantation site. The cortical tissue and 2 layers of corpus callosum fibers above the implantation site were aspirated. Buffered ACSF was applied throughout the aspiration to prevent desiccation of the tissue. A GRIN lens (1 mm diameter, 4.3 mm length, 0.4 pitch, 0.50 numerical aperture, Grintech) was stereotaxically lowered at coordinates -1.75 AP, 1.75 ML, 1.35–1.40 DV (in mm from Bregma) and fixed to the skull surface with ultraviolet-light curable glue (Loctite 4305) and further fixed with dental adhesive (Metabond, Sun Medical) and dental acrylic cement (Simplex Rapid, Kemdent). A metal head bar was attached to the cranium using dental acrylic cement for head-fixing the animal during the microscope mounting.

If the GcaMP6f expression was visible in the implanted mouse, 4 weeks later the animals were anesthetized for the purpose of attaching a baseplate for the microscope above the top of the GRIN lens. The baseplate was cemented into place and the miniscope was unlocked and detached from the baseplate.

#### Behavioral learning task

The mice performed a rewarded spatial navigation task on a round-shaped maze (cheeseboard maze; Dupret et al., 2010). The 120 cm diameter cheeseboard had a total of 177 evenly spaced wells. The rewarded wells were baited with ∼100 μL of condensed milk mixed 1:1 with water.

For the first three days, the mice foraged for rewards baited in randomly selected wells. A different, random set of wells was baited in each trial. Next, we performed a spatial learning task. The mice had to learn two locations with baited wells. The baited wells had fixed locations that were at least 40 cm apart, chosen pseudo randomly for each mouse. Mice started the trial in one of the three locations on the maze: south, east or west. The maze was rotated and wiped with a disinfectant (Dettol) in-between the trials to discourage the use of intra-maze cues. Landmarks of black and white cues were installed on the walls surrounding the maze. The trials were terminated once the mice ate both rewards or after 300 s. Each learning day consisted of 8 trials with 2–4-minute-long breaks between the trials. To minimize the effects of the novel environment and task structure, we analyzed the neural activity on the first learning day after one of the previous reward locations was moved. After the 5-day-long learning, the memory retention was tested on the next day in a 4 to 5-minute-long unbaited trial. Following the learning and testing of the memory for the first set of locations, we translocated one of the reward locations. The new location was a pseudo-randomly chosen to be at least 40 cm away from the current and previous reward locations. The learning of the new set of locations was performed over two days and tested in an unbaited trial as described above.

The trials were recorded with an overhead webcam video camera. The video was recorded at 24 Hz frame rate. The mice body location was tracked with DeepLabCut software (Mathis et al., 2018), a custom-written software was written to map the mouse coordinates to relative location on the maze. The extracted tracks were smoothed with a Gaussian kernel. Periods of running were identified when velocity of the mouse smoothed with a moving average 0.5 s window exceeded 4 cm/s.

#### Calcium imaging

CaImAn software was used to motion-correct any movements between the calcium imaging frames, identify the cells and extract their fluorescence signal from the video recordings (Giovannucci et al., 2019). The method for the cell and signal detection is based on constrained non-negative matrix factorization (CNMF-E; Pnevmatikakis et al., 2016). CaImAn extracted background-subtracted calcium fluorescence values and the deconvolved the signal. The deconvolved signal can be interpreted as a scaled probability of a neuron being active. The calcium imaging videos recorded in the same-day trials were concatenated and motion-corrected to a common template frame. Signal extraction and further processing was performed on the resulting long video, allowing to detect the cells and signal present across the trials. To improve the computational performance, the video frames were cropped to a rectangle containing the regions of interest and the video width and height was downsampled by a factor of 2.

The identified putative cells were automatically filtered using CaImAn. The results were visually inspected and the filtering parameters adjusted to exclude non-cell like shapes and traces from the filtered components. The criteria used for the filtering included a threshold for signal to noise ratio of the trace, the minimum and maximum size of the component‘s spatial footprint, threshold for consistency of the spatial footprint at different times of the component‘s activation, and a threshold for component‘s resemblance to a neuronal soma as evaluated by a convolutional neural network provided with CaImAn software.

The deconvolved trace was time binned, averaging the values in 200 ms bins. A calcium event was detected whenever the cell‘s deconvolved signal crossed 20% of its day-maximum value. A cell was classified as reactivated if it had at least one calcium event when the mouse was at the reward location and it had a calcium event during running preceding the reward. Activity peaks were quantified by their area-under-curve (AUC) which was calculated as a convolution of the preprocessed calcium signal with a 2-s-long flat kernel. The preprocessing of calcium signal subtracted the cell’s median value and truncated the values below 0, so that only the above-median calcium signal is integrated in the AUC calculation.

#### Place cell detection and analysis

To assess how spatial locations modulated activity of a cell, we considered periods of running as described in the ‘Behavioural learning task’ section and calculated place maps — mean neural activity per spatial bin. The total activity inside 6 x 6 cm bins was summed from the smoothed deconvolved signal. The mean neural activity in the spatial bin was then calculated as a ratio of the total activity to the total occupancy in the bin after both maps were smoothed across the space using a 2D Gaussian kernel with σ = 12 cm. The place map was filtered to include spatial bins with total occupancy that exceeded 1 s (5 time bins, thresholded on unsmoothed total occupancy).

Spatial information of a cell‘s activity was calculated using the place map values. Spatial information (Markus et al., 1994) was defined as:

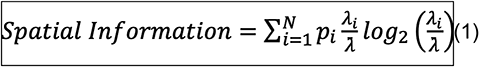

where 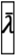 represents the mean value of the neural signal, *p_i_* represents probability of the occupancy of the i-th bin, and *λ_i_* represents its mean neural activity. Dividing by 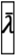 ensures the metric is independent of the cell‘s average activity. The units of spatial information calculated on calcium fluorescence can be reported as bits per action potential (Climer and Dombeck, 2021). However, because the actual action potentials were not measured, we report them as arbitrary units.

Spatial information was compared to the value expected by chance. The chance level was calculated by circularly shifting the activity with regards to the actual location. For each cell, the activity was circularly shifted within the trial by a time offset chosen randomly (minimum offset 10 s for baited and 20 s for unbaited trials). If the cell‘s spatial information exceeded 95% values calculated on 1000 random shifts of its activity, it was defined as a place cell.

A limited number of neuronal responses sampled per spatial bin can lead to an upward bias in the estimated spatial information (Treves and Panzeri, 1995). To correct for this bias, we subtracted its estimated value from the estimated spatial information. The bias was estimated as the mean spatial information from the time-shifting procedure used for place cells detection. This procedure does not require binning the neuronal responses from the calcium imaging as required by analytical estimation (Panzeri et al., 2007), and has been used previously to estimate mutual information bias (Akrami et al., 2018).

#### Statistical testing of in vivo results

To compare the activity peaks between the reactivated and non-reactivated cells throughout the day, we used permutation tests for repeated measures ANOVA (Figure 8E, Figure 8-S1A,B, Figure 8-S2). The ANOVA modelled fixed effects of trial ordinal and reactivation and random effects of within animal-session factors. To compare cells grouped by their first activation trial, we used permutation t-tests (Figure 8F, Figure 8-S1C). Multiple comparisons were corrected with Benjamini-Hochberg method, and the permutations were restricted to within animal-session permutations. Both effects were computed using ‘permuco‘ R package. Statistical analysis was performed in R version 3.6.3.

#### Data availability

Experimental data and code are available at https://github.com/ … [To be completed].

---

**Figure 8-S1.**
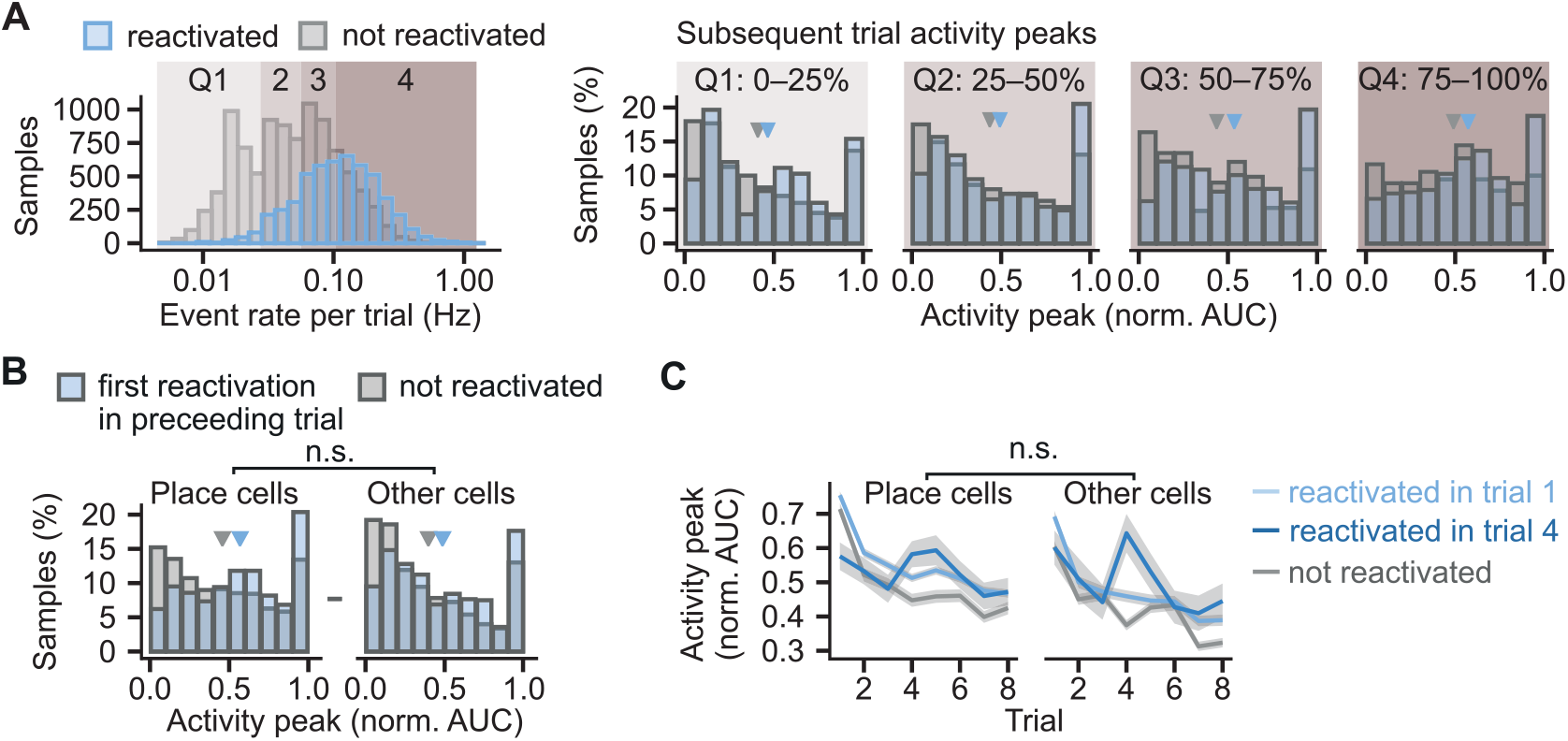
Similar effect of the reactivation on the activity peaks in different cell groups. (A), The activity peaks were larger in the reactivated than in the non-reactivated throughout the day cells regardless of cell event rates. The comparison was performed on four cell groups defined by the quartile ranges (Q1–4) of event rate distribution. The leftmost histogram plots the trial event rates per cell. The four histograms to the right compare within each quartile activity peaks of the same cells in the subsequent trial. Triangles mark mean values. Permutation test for repeated measures ANOVA: effect of reactivation: F(1, 16) = 6.33, p = 0.02; effect of quartile: F(3, 48) = 6.76 , p = 0.001. n = 6,447 cell trials from 1,404 cells. (B), Histogram of normalized activity peaks in place cells and other cells compared between the cells reactivated for the first time in the preceding trial and the cells not reactivated in any of the trials. Triangles mark mean values. Permutation test for repeated measures ANOVA: non-significant interaction between cell type (place cell vs other cell) and reactivation (F(1,16) = 0.16, p = 0.69, n = 6,496 cell trials from 1,404 cells). (C), Activity peaks as a function of trial compared between place cells and other cells. Cells whose first reactivation happened in the same trial are grouped together. Ribbons extend +-1 SEM. The difference in the activity peaks was not-significant between place cells in other cells (p = 0.88, n = 6,496 cell trials from 1,404 cells, permutation t-tests with Benjamini-Hochberg correction for multiple comparisons and between-animal random effects. *p < 0.05).

**Figure 8-S2.**
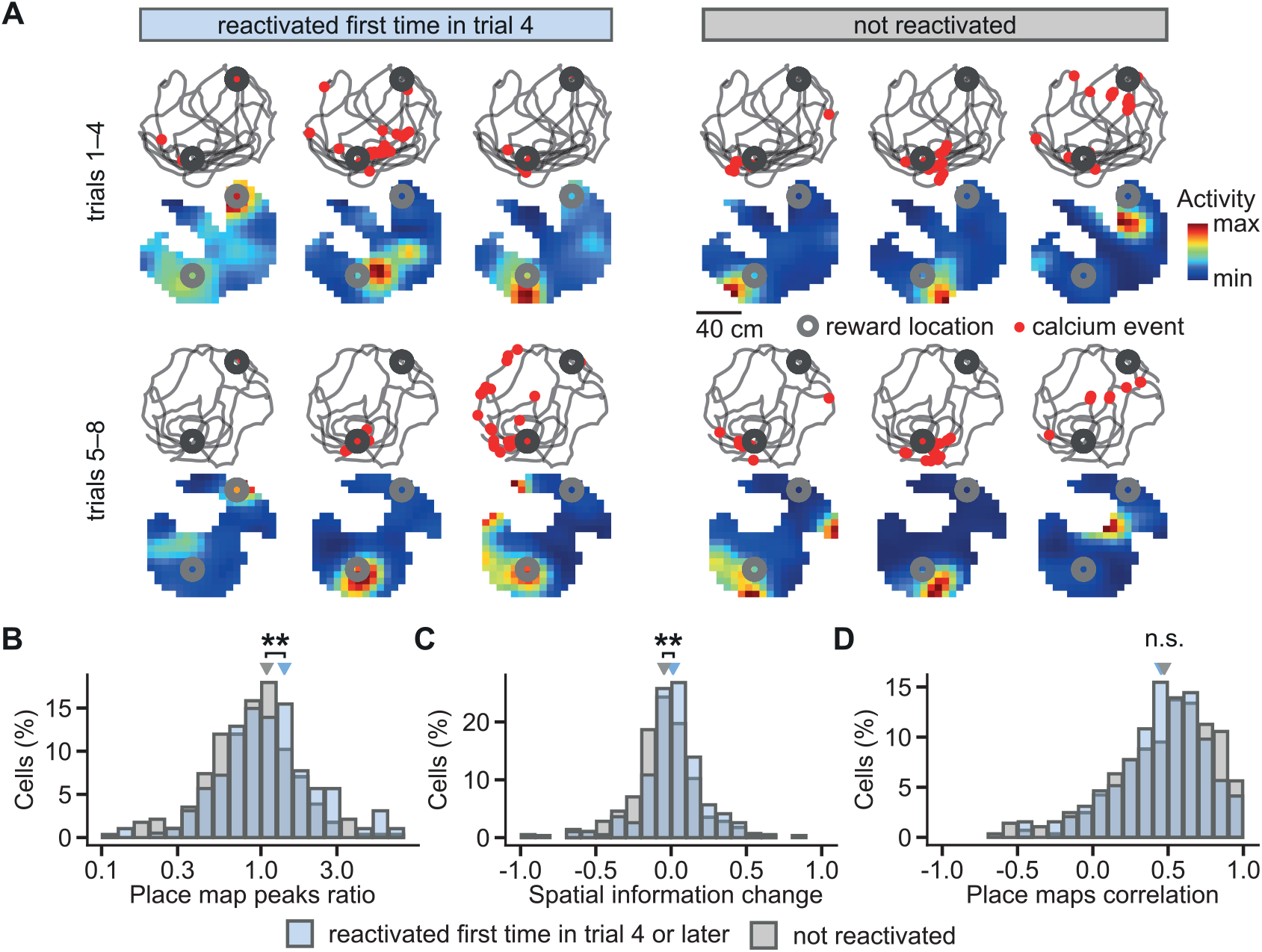
Place cells during learning in reactivated and non-reactivated neurons. (A), Examples of CA1 place cells and changes in their activity from early (trials 1–4) to late learning trials (trials 5–8) after the cells were reactivated at the reward location in trial 4. Locations of calcium events marked with a red dot are overlaid over mouse movement paths; place maps are shown below. Reward locations are marked with grey circles. (B), Histogram of the ratios between place map peaks in the late over early learning trials. The ratios are compared between the cells reactivated for the first time in the trial 4 or later and the cells not reactivated at any of the trials. Triangles mark mean values. (C), As in (B) but for change in spatial information of place cells from early to late trials. (D), As in (B) but for correlation between place maps for early and late trials. Permutation tests for repeated measures ANOVA. **p < 0.01.

## Notes

### Competing Interest Statement

The authors have declared no competing interest.

### Summary of Updates

Revision merely to correct errors to PDF conversion.

